# Sample-multiplexed FACS-preprocessing of PBMCs enables scalable scRNA-seq without compromising transcriptomic or cellular integrity

**DOI:** 10.1101/2025.11.06.687014

**Authors:** Mohammad Mokhtari, Timothy J.S. Ramnarine, Antonia Eicher, Alexander Braunsperger, Philipp Baumert, Christine Wolf, Görkem Durmaz, Veronika Pfaffenstaller, Arek Kendirli, Martin Kerschensteiner, Martin Schönfelder, Min Ae Lee-Kirsch, Henning Wackerhage, Simon W. Mages, Johanna Klughammer

**Author notes:** Equal contribution (Mohammad Mokhtari, Timothy J.S. Ramnarine). Joint supervision (Simon W. Mages, Johanna Klughammer). Corresponding authors: Johanna Klughammer, Gene Center and Department of Biochemistry, Ludwig-Maximilians-Universität München, Feodor-Lynen-Straße 25, 81377 Munich, Germany., Simon W. Mages, Gene Center and Department of Biochemistry, Ludwig-Maximilians-Universität München, Feodor-Lynen-Straße 25, 81377 Munich, Germany.

## Abstract

Efficient preprocessing of peripheral blood mononuclear cells (PBMCs) for single-cell RNA-Sequencing (scRNA-seq) is crucial to ensure high sample throughput while maintaining sample integrity. In particular, when enrichment of rare immune cell populations is necessary to enable their representative profiling among more common PBMCs, sample preprocessing may become a detrimental bottleneck. Here, we present an optimized fluorescence-activated cell sorting (FACS)–based preprocessing workflow designed to enrich rare immune cells while conserving overall PBMC composition. The protocol integrates dead cell removal, targeted rare cell enrichment, channel splitting, and hash-based sample multiplexing together with a new powerful yet lightweight demultiplexing tool (YAHD), improving throughput and cell yield, reducing batch effects, and preserving biological context. Validation across cryopreserved human PBMCs obtained from different scientifically relevant sources (clinical routine and laboratory setting) demonstrated improved sample viability and representation of rare subsets in the final scRNA-seq data. Thorough transcriptomic assessment confirmed non-concerning levels of stress induction and T cell activation as well as low technical variability, removing concerns around FACS-processing, cross-donor multiplexing and channel splitting. The presented approach enables scalable and biologically faithful PBMC preprocessing for scRNA-seq, advancing the study of immune heterogeneity in health and disease.

## Background

Human peripheral blood mononuclear cells (PBMCs) are a widely used resource in immunological and clinical research. They are obtainable in a minimally invasive manner, easily accessible, relatively inexpensive to isolate, easy to handle and preserve, and capture both innate and adaptive immune compartments within a single blood draw (1–4). Millions of cells can be obtained from an individual sample, making PBMCs suitable for studying immune dynamics in diverse contexts ranging from dynamic physiological changes over immune disorders to malignant diseases (5–9). However, PBMC isolates are often not directly suitable as input to sensitive transcriptomic profiling such as scRNA-seq due to erythrocyte contamination, low sample viability, and low frequencies of certain cell types of interest. Especially in the clinical setting, where samples often cannot be immediately processed, overall sample quality may additionally suffer as opposed to the controlled laboratory setting. Cryopreservation of PBMCs, which is often required for logistical reasons, may further affect sample quality and transcriptomic integrity. Preprocessing of PBMC isolates prior to scRNA-seq is therefore aimed at optimizing the input sample such that resources are spent on cells of interest and not wasted on dead cells, debris, or highly frequent cells with limited transcriptomic information such as erythrocytes. At the same time, the overall cellular composition should be maintained as faithfully as possible while also capturing rare, yet important subsets such as regulatory T cells, innate lymphoid cells, and dendritic cell types (10–13) in sufficiently high numbers to enable their robust characterization. Another important consideration during sample preprocessing is to limit stress that cells experience, rendering protocol gentleness, efficiency and scalability increasingly important as larger numbers of samples are processed.

Sample multiplexing through antibody- or lipid-based hashing has emerged as a popular strategy in scRNA-Seq to process large numbers of samples, especially when samples are pooled before preprocessing (14–16). However, many popular enrichment or dead cell removal approaches such as magnetic cell sorting, (17,18) are less compatible with a multiplexed setup because they typically require temperatures above 4℃, at which the hashing tags attachment to the cells becomes unstable and concerns about immune activation (alloreactivity) between immune cells of different donors may increase (19,20). Recent systems employing levitation-based processing (e.g. Levitas Bio: LeviCell) in single-cell studies (21,22) have been updated to include cooling, parallel sample capabilities and limited antibody-based enrichment options (e.g. Levitas Bio: LeviSelect) but the latter is currently restricted to common immune cell subtypes such as T cells, B cells, NK cells, and monocytes, rather than rare or specialized cell populations (manufacturer documentation). Other solutions (e.g. Parse Biosciences: Evercode or 10x Genomics: Flex kit) can achieve near-complete hashtagging efficiency but still require separate sample processing for rare cell enrichment (23,24). In contrast, fluorescence-activated cell sorting (FACS) can be performed at 4℃. In principle, FACS allows simultaneous dead cell removal, rare cell type enrichment, and - with appropriate gating strategies-preservation of proportions among the more frequent cell types (25,26). These properties make FACS-based preprocessing popular among scRNA-seq studies (27–32). However, despite its common use no standardized and validated FACS-based PBMC preprocessing protocol for high-throughput sample-multiplexed scRNA-seq experiments is available, frequently leading to concerns around potential biases in the resulting data (33–35).

Here, we present an optimized and validated preprocessing protocol for scalable sample-multiplexed, FACS preprocessed scRNA-seq with detailed documentation available on protocols.io (https://dx.doi.org/10.17504/protocols.io.j8nlk82m5l5r/v1) and the fast, flexible and Python-based demultiplexing tool YAHD (“Yet Another Hash Demultiplexer”, github.com/simonwm/yahd), that delivers robust demultiplexing performance also in cases when the standard 10x Genomics Cell Ranger (36) integrated demultiplexing fails.

We use FACS in combination with antibody-based sample multiplexing because this approach is compatible with high sample throughput while providing optimal control over key factors such as temperature, cell viability, relative cell proportion, and labeling specificity, which is critical for minimizing technical artifacts in multiplexed single-cell workflows (25,26). Our protocol integrates dead cell removal and rare cell enrichment directly within multiplexed, antibody hashtagged PBMC samples, thereby efficiently enhancing representation of low-frequency populations without distorting the broader cellular composition and, through the removal of dead cells, mitigating dead cell and ambient RNA contamination (38,39). Sample multiplexing through cell hashing not only streamlines processing by increasing throughput, but also reduces batch effects, multiplet rates and facilitates accurate doublet detection, including homotypic doublets which are otherwise hard to detect (16). Combined with channel splitting (processing the same cell suspension on multiple 10x channels), multiplexing allows more efficient use of channel capacity, improved recovery across samples and technical safeguards against sample loss due to channel failure.

We thoroughly validate the approach across 16 scRNA-seq runs and a total of 112 samples including 26 control samples in the clinical (8 runs, 64 total samples, 10 healthy control samples) as well as laboratory-based (8 runs, 48 total samples, 16 baseline control samples) setting, while disentangling connections between sample viability, cellular stress and sequencing depth with important implications on sample processing and data analysis. We confirm improvement of input sample quality in the final scRNA-seq data and assess cellular stress, activation, and alloreactivity signatures as well as technical variability, while investigating and correcting surprising effects of sequencing depth variability which becomes especially relevant at shallow sequencing depths that are recommended for cost-optimization in scRNA-seq studies (40).

## Results and Discussion

To optimize data quality, we implemented a stepwise strategy with the following preprocessing steps prior to scRNA-seq library preparation using the 10x 5’ v2 protocol: sample-multiplexing, dead cell removal, rare cell enrichment, and channel splitting. Our strategy was applied in two contexts: (i) PBMCs collected from healthy participants under controlled laboratory conditions with optimal sample treatment (“Lab-sourced” dataset), and (ii) PBMCs collected from patients in more variable clinical settings where samples typically wait longer until they are processed (“Clinical” dataset) (**Fig. 1a**).

**Figure 1.**
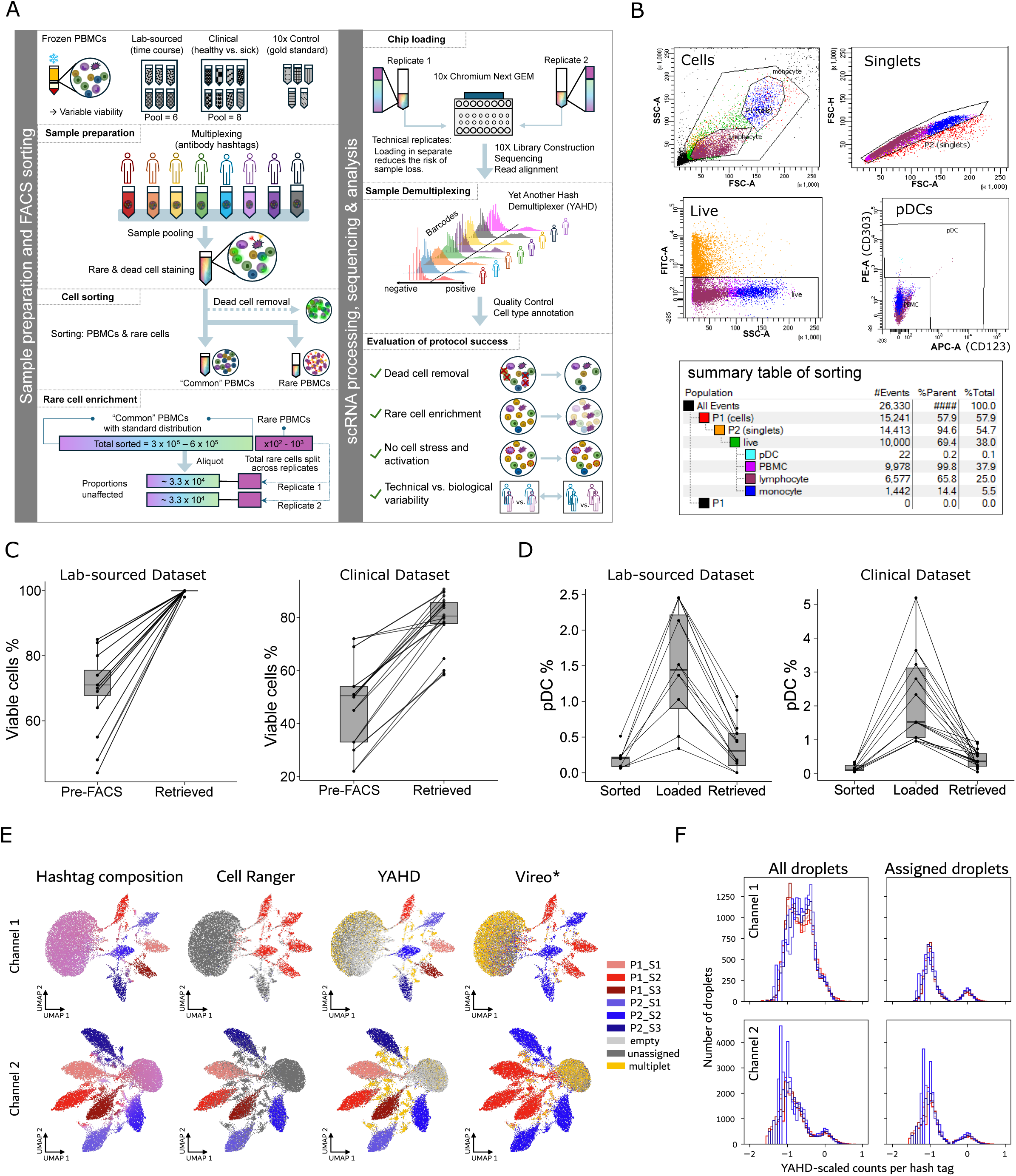
Sample-multiplexed, FACS-based preprocessing for scRNA-seq. (A) Overview of the preprocessing workflow comprising five major steps: sample multiplexing, dead cell removal, rare cell enrichment, and channel splitting, and the analytical approach for demultiplexing and workflow validation. (B) Representative FACS gating strategy illustrating key cell filtering and sorting steps performed on multiplexed PBMC pools prior to 10x chip loading. Gating was applied sequentially to identify living cells, exclude multiplets, and remove dead cells based on live/dead staining. Subsequent gates were used to enrich for pDCs and to quantify the fraction of cells retained at each step. (C) Boxplots showing the percentage of living cells before FACS (“Pre-FACS”) and after recovery in scRNA-seq analysis (“Retrieved”) for the Lab sourced and Clinical datasets. Post-analysis viability was assessed by defining live cells as those with <25% mitochondrial UMI counts. (D) Box plots showing the proportion of plasmacytoid dendritic cells (pDCs) at three consecutive stages of the workflow: after FACS sorting, at the loading stage when pDCs were pooled with “other” PBMCs for 10x processing, and among retrieved cell-type annotated cells in the final scRNA-seq data. (E,F). Hash tag demultiplexing for a partially clogged 10x channel (channel 1) and a non-clogged channel (channel 2) from the same cell suspension. (E) UMAP embedding of all droplets with at least 100 expressed genes and at least 10 hash tag counts colored by fractional composition of the hash tag counts (left), the hash tag demultiplexing result by Cell Ranger multi (center left), the hash tag demultiplexing result by YAHD (center right), and the genetic demultiplexing result by Vireo (right). *As the six samples correspond to only two donors, the Vireo result is shown by one representative color for each donor. (F) Histograms of the YAHD-scaled hash tag counts per cell for all droplets shown in (E) (left) or only for the subset of droplets which were assigned a single sample by YAHD, i.e. without the categories “empty”, “unassigned”, and “multiplet” (right).

In a first step, samples were multiplexed in pools of six (Lab-sourced) or eight (Clinical) through antibody-based hashing, with each pool containing samples from two (Lab-sourced) or up to eight (Clinical) unrelated donors including two base-line controls (Lab-sourced) or at least one immunologically healthy control (Clinical). All following steps, including FACS, were then performed on the sample pools and finally each pool was processed on two 10x channels to produce the desired number of 3000-4000 cells per sample (**Fig. 1a,b**). For protocol validation, only base-line or healthy controls were used to avoid interfering biological signals and to ensure comparability to data published by 10x Genomics that we included as gold standard.

For dead cell removal and enrichment of rare pDCs (0.1-0.5% of PBMCs), we established a flow cytometric gating strategy that sequentially excluded debris, doublets, and dead cells, while sorting pDCs and other PBMCs into different tubes (**Fig. 1b**). Live cells were identified by exclusion of high Sytox Green–positive events, and pDCs were defined as CD123^high and CD303^high cells (41). All other events were notated as “other PBMCs” encompassing all monocytes and lymphocytes in their native proportions. We validated the effectiveness of dead cell removal, by comparing cell viability before sorting and in the scRNA-seq data defining dead cells as those with >25% mitochondrial counts. In the Clinical dataset, mean viability increased from 47.7% to 78.9%, an absolute gain of 31.2 percentage points (∼ 66% relative increase), while in the Lab-sourced dataset, mean viability increased from 69.7% to 99.9%, an absolute gain of 30.2 percentage points (∼43% relative increase), demonstrating consistent improvement across datasets (**Fig. 1c**).

Rare cell enrichment was confirmed by tracking pDC proportion at three stages: (i) the FACS cell sorting output, (ii) the proportion of pDCs loaded alongside “other” PBMCs, and (iii) the number of annotated pDCs retrieved after sequencing and annotation (**Fig. 1d**). At the loading stage, the pDC fraction was several-fold higher than the initial FACS proportion (Clinical mean: from 0.16% to 2.20%; Lab-sourced, mean: from 0.20% to 1.47%), reflecting the fact that we deliberately pooled pDCs collected from many hundreds of thousands of sorted cells into ∼33k “other” PBMCs, thereby concentrating the rare population for capture. A reduction was then observed from loaded to retrieved fractions (Clinical mean: from 2.20% to 0.42%; Lab-sourced, mean: from 1.47% to 0.35%), likely due to losses during pellet washing and the inherent capture efficiency of scRNA-seq. Crucially, the retrieved proportions remained enriched relative to the native proportion represented in the FACS sorting report (Clinical mean: from 0.16% to 0.42%, +0.26 points, 163% relative increase; Lab-sourced, mean: from 0.20% to 0.35%, +0.15 points, 75% relative increase), confirming robust recovery of this rare population (**Fig. 1d**).

Within this workflow, upfront sample-multiplexing is a well-established, cost-saving strategy that reduces batch effects, maintains cell type frequencies, and saves time by enabling simultaneous processing of multiple samples. It additionally increases the computational identifiability of doublets which allows us to increase the target number of recovered cells per 10x channel from 5k to 20k at a residual doublet rate of <= 4% (16). Importantly, sequencing reads are still wasted on doublets (16% at 20k recovered cells) which increases the sequencing cost per non-doublet cell and thus limits the advantages of increasing the number of targeted cells beyond 20k. We processed each multiplex on two 10x channels (channel splitting), allowing us to obtain the desired number of cells per sample, while generating technical replicates, and creating a safeguard against complete sample loss in case of channel failure (**Fig. 1a,b**).

While using the 10x recommended demultiplexing strategy included in 10x Genomics Cell Ranger multi(36), for some runs we noticed complete or partial failure to resolve hash tags in one channel but not the other indicating a problem with the demultiplexing approach rather than the data. We traced the problem to unwarranted flagging of hash tags as contaminating and low signal-to-noise ratios. To resolve these issues, we developed “Yet Another Hashing Demultiplexer” (YAHD), a light-weight, fast, flexible, and robust Python tool that successfully resolved the problematic channels. We confirmed YAHD-based donor assignments by comparing them with genetic demultiplexing results from Vireo (37), Cell Ranger, and plain hashtag compositions (**Fig. 1e**). Plain hashtag composition i.e. coloring droplets by their relative “expression” of the six different hash tags clearly showed the expected distribution for both channels, confirming rather a problem with computational assignment than the data. By design, Vireo was only able to assign the two donors but not distinguish samples from the same donor. All tested methods were able to consistently assign cells in channel 2, but Cell Ranger was unable to correctly assign samples in channel 1 assigning all cells to a single sample or leaving them unassigned, whereas YAHD and Vireo consistently assigned samples and donors, respectively (**Fig. 1e**). Assessing the distribution of YAHD-scaled counts per cell demonstrated the absence of clear signal separation between hash tag positive and negative cells before YAHD-assignment in channel 1, while clear signals were always observed in channel 2 and in both channels after YAHD-assignment, confirming that YAHD robustly demultiplexes even suboptimal initial hashtag signal distributions (**Fig. 1f**).

Despite the clear advantages of sample multiplexing, it raises the possibility that the required additional preprocessing may induce transcriptional stress responses. Another potential concern when pooling PBMC samples from different donors is non-self-reactivity, in which HLA mismatches could trigger T cell activation and related transcriptional responses. Although recent benchmarking suggests that this risk is minimal under standard scRNA-seq preparation conditions (20), we sought to confirm this explicitly (**Fig. 2a,b**). To rigorously test stress and non-self-reactivity signatures in our presented protocol, we compared stress and non-self-reactivity expression signatures in the Lab-sourced and Clinical datasets against multiple publicly available PBMC best-practice (“control”) datasets from 10x Genomics, using concordant cell type annotations and QC filtering across all datasets (**Fig. 2b, Supplemental Fig. S1a**)

**Figure 2.**
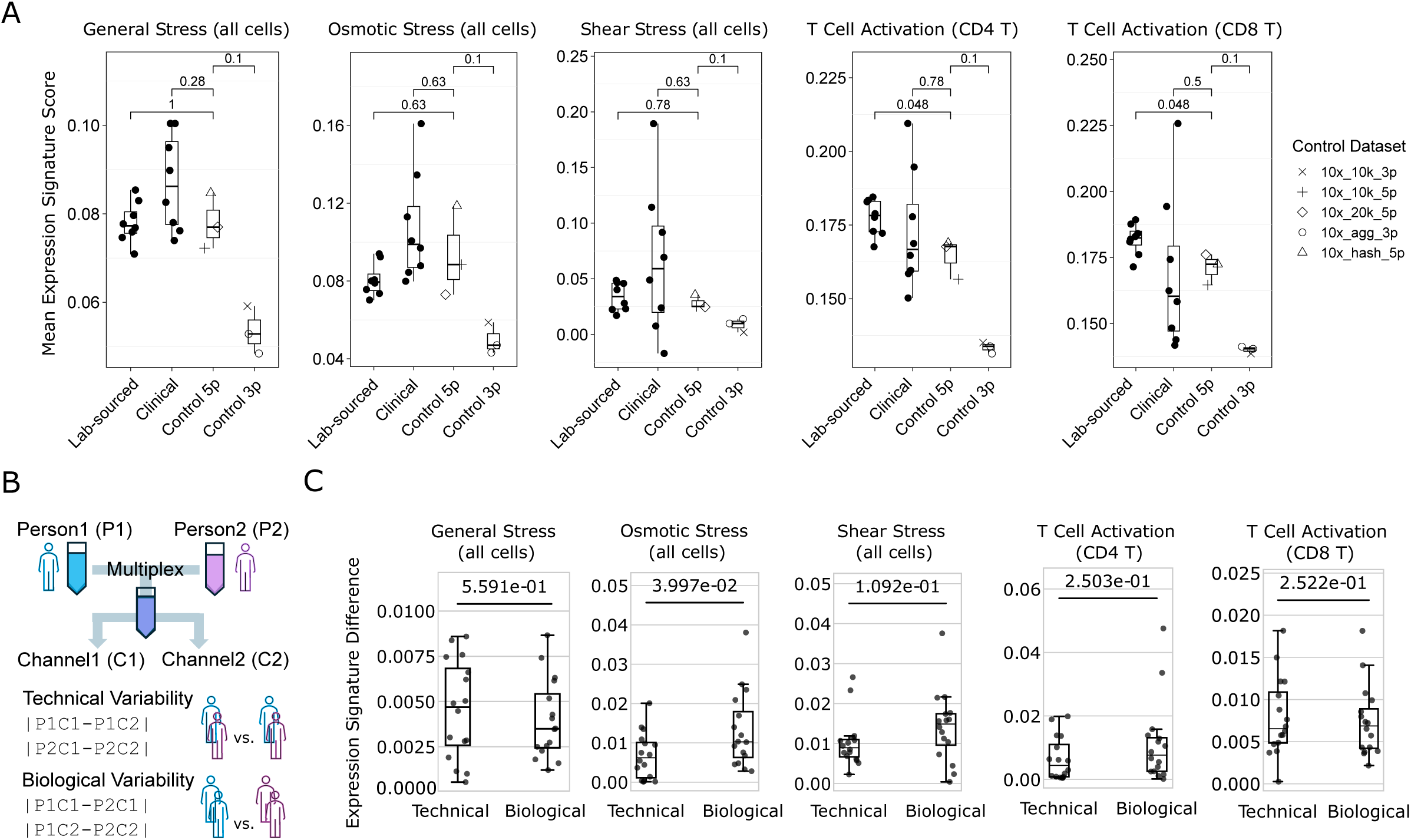
Assessment of cellular transcriptional stress and activation levels. (A) Expression signature analysis of general stress, osmotic stress, fluid shear stress, and T-cell activation gene sets in CD4⁺ and CD8⁺ T cells across Lab-sourced, Clinical, and publicly available 10x control datasets (5′ and 3′ chemistries). (B) Schematic of the experimental design for assessing biological versus technical variability in the Lab-sourced dataset. Biological variability was defined as inter-individual differences within the same 10x chip channel, while technical variability corresponded to channel-to-channel differences for the same biological sample processed across two 10x chip channels. (C) Comparison of general stress, osmotic stress, fluid shear stress, and T-cell activation signature score differences between biological and technical replicates in the Lab-sourced dataset.

Upon QC we observed quite drastic differences in counts per cell between datasets (**Supplemental Fig. S1b top**), likely owing to differences in sequencing depth with our datasets being on the lower end due to cost optimization for cells over sequencing depth as previously recommended (40). To check whether differences in counts per cell may affect expression signature scoring, we performed an in-silico downsampling experiment with one of the 10x control samples and found that large gene-sets (>2000 genes) such as the “general stress” gene set (GO:0033554) achieved increasingly lower scores with increasing degrees of downsampling, while scores for small gene sets (<100 genes) or sets of randomly selected genes appeared to be unaffected by downsampling (**Supplemental Fig. S1c**). Thus, to ensure the comparability of expression signature scores, we downsampled all datasets to a common virtual sequencing depth achieving similar counts per cell distributions across libraries (**Supplemental Fig. S1b bottom**). While in the Lab-sourced dataset, downsampling removed the artefactual positive correlation between library-wise general stress signature scores and mean counts per cell (R=0.73 → R=-0.09), in the Clinical dataset improvement was marginal (R=0.89 → R=0.70) (**Supplemental Fig. S1d**), indicating additional factors driving the spurious correlation. Indeed, after downsampling, both datasets showed the biologically expected negative correlation between general stress signature scores and sample viability (Lab-sourced: R=-0.66, Clinical: R= -0.48) (**Supplemental Fig. S1d**), but while in the Lab-sourced dataset sample viability was positively correlated to mean counts per cells (R=0.41), in the Clinical dataset it was negatively correlated (R=-0.45) (**Supplemental Fig. S1e top**), presenting sample viability as a biological explanation for the persistent correlation between general stress signature scores and mean counts per cell in the Clinical dataset. Additionally, we observed that samples of both datasets with a viability below ∼55% seemed to follow the negative viability vs. mean counts per cells correlation trajectory, while those with a viability greater than ∼55% followed positive correlation trajectory, indicating a dataset agnostic phenomenon as root cause (**Supplemental Fig. S1e top**). Indeed, we found that in both datasets sample viability was positively correlated with the number of cells profiled (Lab-sourced: R=0.23, Clinical: R= 0.70) (**Supplemental Fig. S1e bottom**), providing a direct link between sample viability and mean counts per cell: In samples with low viability the same number of sequencing reads is distributed among less cells because cells from lower viability samples (<∼55% viability), which are more frequent in the Clinical dataset **(Fig. 1b**), continue dying after initially loading the same number of live cells.

Having confirmed the validity of our downsampling strategy to account for sequencing depth differences, we evaluated stress-related and immune-activation-related expression signatures across Lab-sourced, Clinical, and control datasets. Signatures included general stress (GO:0033554), osmotic stress (GO:0071470), fluid shear stress (GO:0034405), and T-cell activation (GO:0042110), measured in CD4+ and CD8+ T cells separately (**Fig. 2a**). While the Clinical dataset displayed higher variability between libraries resulting from higher variability in viability **(Fig. 2a, Supplemental Fig. S1e**), no significant differences were found between Lab-sourced/Clinical and 5’ 10x control datasets. However, 3’ 10x control datasets showed significantly lower signature scores compared to all other datasets, indicating important differences between 10x chemistries (**Fig. 2a**). Overall, these results indicate that our protocol does not impose additional stress or trigger non-self-reactivity(42,43), supporting its reliability in both clinical and lab-based contexts.

Finally, we wanted to assess the level of variability introduced by our channel-splitting strategy (i.e. processing the same pool of samples on more than one 10x channel to achieve sufficiently high cell numbers for each sample). In the Lab-sourced dataset, the experimental design enabled direct comparisons between biological replicates (samples from two different participants per pool) and technical replicates (two 10x channels used to process each pool). Specifically, biological variability was defined as inter-individual differences within the same 10x channel, while technical variability reflected channel-to-channel differences within the same biological sample (**Fig. 2b**). Stress and T cell activation expression signature differences were low overall and showed that both sources of variability were comparable, with a slight trend toward higher biological variability (**Fig. 2c**). This indicates that the technical variation between 10x channels was never meaningfully greater than biological variation between biological replicates (base-line control samples) and thus observed differences in measured stress and T-cell activation signatures primarily reflect genuine biology rather than protocol-related artifacts.

Our results show that integrating sample-multiplexing, FACS-based dead cell removal and rare cell enrichment, and channel splitting provides a robust and scalable workflow for single-cell RNA-seq of PBMCs without introducing stress or cell activation related biases. This strategy improves sensitivity for rare populations, preserves in vivo cell type proportions, minimizes technical artifacts and is applicable in the controlled laboratory setting as well as the more variable clinical setting. Comparing data from both setups, together spanning a wide range of sample viability (20-85%), additionally revealed an important turning point at ∼55% sample viability below which cells continue dying during processing, leading to reduced cell recovery if not counteracted by loading proportionally more cells. Data-analytically, we demonstrate the necessity to take into account sequencing depth differences between libraries when calculating and comparing gene expression signatures and we introduce YAHD as a light-weight demultiplexing tool that delivers robust demultiplexing performance also in non-trivial cases.

Overall, the methodological robustness and scalability of the protocol together with our thorough validation and analysis provide a strong foundation for future studies aiming to capture with high fidelity common and rare cell populations also in challenging samples, ultimately enabling more accurate and efficient biological discovery.

## Methods

### Study design and sample donors

Samples were obtained from two distinct sources, referred to as “Lab-sourced” and “Clinical” in the context of two distinct biomedical research projects, studying interferonopathies (Clinical) and the effect of exercise on the immune system (Lab-sourced). The protocol and demultiplexing tool presented in this work were developed to enable efficient scRNA-seq data generation for those two (and future) research projects. In this study, the resulting scRNA-seq data were exclusively analyzed for the purpose of assessing the performance of our protocol and analysis were only performed on healthy control (Clinical) or baseline-control (Lab-sourced) samples to avoid any interference of biological signals.

Written informed consent was obtained from all participants or their legal guardians. The Clinical study protocol was approved by the ethics committees of the Medical Faculty, Technische Universität Dresden, Charité-Universitätsmedizin Berlin, and the Lab-sourced study protocol was approved by the ethics committee of Technical University of Munich and both were conducted in accordance with the Declaration of Helsinki.

### Sample Acquisition: Lab-sourced samples

Blood samples were obtained from healthy young adults (23-28 years old) participating in an exercise study. Participants enrolled into the study and completed a spiroergometric test (Cortex Metalyzer 3b, Cortex, Leipzig, Germany) to assess individual maximal aerobic capacity (VO_2_max) during a ramp test on a cycling ergometer (Lode Excalibur Sport, Lode, Groningen, Netherlands) until maximal exhaustion. On a separate day, venous blood was drawn at rest after a 12 h overnight fast (baseline-control time point), after 60 minutes of moderate intensity continuous cycling (60% of individual VO_2_max), and at last, 30 minutes post cycling. In this study, only the baseline-control time point was incorporated. Participants underwent strict inclusion-exclusion screening to control for immunological background, medication use, alcohol consumption, and smoking status. At the study site, 9 ml of blood was drawn via multifly safety needle 21 G into K₂EDTA tubes to prevent coagulation. Samples were immediately processed on-site to isolate PBMCs, minimizing any delay between blood draw and processing. Baseline control samples were collected at rest, following approximately one hour of rest after arrival in the laboratory.

### Sample Acquisition: Clinical samples

Control blood samples were obtained from healthy donors (5-35 years old) admitted to the University Hospital Dresden for non-immune-related conditions (e.g. orthopedic check-ups). Donors were screened to exclude acute or chronic infections. Unlike Lab-sourced samples, Clinical samples were subject to the hospital’s clinical workflow, resulting in a delay of several hours between blood draw and PBMC processing. Blood was collected into K₂EDTA tubes and temporarily stored at room temperature until processing.

### PBMC isolation and cryopreservation

Peripheral blood was diluted in phosphate-buffered saline (PBS) and carefully layered over 9 mL Ficoll (Biocoll) for density gradient centrifugation to isolate the buffy coat containing peripheral blood mononuclear cells (PBMCs). The PBMC layer was collected and washed 2– 3 times in PBS by sequential dilution and centrifugation. After the final wash, cells were pelleted, resuspended in culture medium, and transferred into cryovials. Freezing medium was added dropwise while gently mixing. Cryovials were placed in a pre-cooled freezing container, stored at −80 °C for 4 hours, and subsequently transferred to liquid nitrogen for long-term storage.

### Sample pre-processing and scRNA-seq

Cryopreserved PBMCs were thawed in a 37 °C water bath and resuspended in 13 mL 5% FBS/PBS (FACS Buffer, FB) to dilute DMSO. After centrifugation (550 x g, 5 min, 4 °C), cell count and viability were assessed using a Countess II Automated Cell Counter (Invitrogen). The cell count and viability at this stage were recorded for downstream analysis. Cells were resuspended in 45 µL FB, blocked with 5 µL Human TruStain FcX (BioLegend) for 10 minutes, and stained with TotalSeq-C anti-human Hashtag antibodies and pDC markers CD123 and CD303 (BioLegend) for 20 min at 4 °C. After incubation, excess hashtag and antibody dye were washed from each sample by adding approximately 10 mL of FB to each sample and centrifuging (550 x g, 5 min, 4 °C). The supernatant was removed and the pellet was resuspended in 0.5-1.0 mL FB. Cell count and viability were again assessed and recorded using a Countess II Automated Cell Counter (Invitrogen). Samples were pooled according to viable cell numbers at this stage such that all samples were equally represented in the pool. The resulting pool of cells was then sorted by FACS to remove dead cells and enrich for pDCs. Sorted pDCs and remaining PBMCs were collected separately and the total sorted counts were recorded.

The sorted pDCs were kept on ice, and 300 µL of 0.1% BSA/PBS was immediately added to the collection tube. Half of this volume was transferred to a new Protein LoBind tube, while the remaining half was left in the original sorting tube to minimize potential cell loss during pipetting. This resulted in two tubes containing equal fractions of the sorted pDCs, which were used as technical replicates.

Sorted PBMCs were then centrifuged (550 x g, 5 min, 4 °C), and the supernatant was removed. The PBMC pellet was resuspended in 100 µL of 0.1% BSA/PBS, and a second cell count and viability assessment was performed using a Countess II Automated Cell Counter (Invitrogen). The cell count and viability at this stage were recorded for downstream analysis. Based on these measurements, the appropriate volume of PBMCs was calculated and added to each tube containing pDCs so that approximately 33,000 PBMCs were supplemented to each technical replicate.

Each technical replicate was then centrifuged a final time (550 x g, 5 min, 4 °C) and the supernatant was carefully removed, leaving ≤ 38.7 µL which is the maximum total load volume for the combined cell suspension and water according to 10x protocol (CG000330 Rev F). This remaining volume was used to resuspend the pellet and was measured by pipetting to determine the exact amount of water to be added to the GEM master mix.

Each technical replicate was loaded onto a separate channel of a 10x Chromium Next GEM Chip K using the Chromium Next GEM Single Cell 5′ Kit v2 (Dual Index) with Feature Barcode technology (CG000330 Rev F). Libraries were sequenced on an Illumina NextSeq 2000 sequencer with 400 million reads per gene expression library and 100 million reads per hash library according to the manufacturer’s protocols.

### 10x gold-standard control datasets

Five publicly available datasets from the 10x Genomics repository were included as control datasets in this study (Table 1):

**Table 1:** Overview of control 10x gold-standard datasets included in this study.

| 10x Dataset | Cryopreserved | Multiplexing | Chemistry | Sample status | Miscellaneous |
| --- | --- | --- | --- | --- | --- |
| Control 10k 5p | NA | - | 5' v2(44) | 1 healthy; male | - |
| Control hash 5p | Yes | Yes. TotalSeq-C antibodies | 5' v2 HT | 1 ALL bone marrow; 1 ALL PBMC; 2 healthy PBMC | Only the healthy samples were used |
| Control 20k 5p | NA | - | 5' v2 HT | 1 healthy;male | - |
| Control 10k 3p | NA | - | 3' v3.1 | 1 healthy;female | - |
| Control agg 3p | Yes | - | 3' v3.1 | 1healthy | 1 chip manual, 1 chip Chromium Connect |

“Control 10k 5p” dataset contains 10 thousand cells from a healthy donor prior to the filterings and was generated with “Chromium Next GEM Single Cell 5’ Reagent Kits v2 (Dual Index)”(44). “Control hash 5p” dataset originally contains samples from 2 healthy donors and 2 donors diagnosed with Acute Lymphocytic Leukemia (ALL). In this study, the cells from ALL donors were filtered out. Samples in this dataset were multiplexed with “TotalSeq™ - C Human Universal Cocktail and a unique cell hashing antibody” and the library was generated with “Single Cell V(D)J Reagent Kits CG000424 (5’ v2 HT)” (45). “Control 20k 5p” is sourced from a healthy male donor and the library is generated with “Chromium Next GEM Single Cell 5’ HT Reagent Kits v2 (CG000421)” and contains 20k cells before the filtering (46). “Control 10k 3p” is sourced from a healthy male donor and the library is generated with “Chromium Single Cell 3’ Reagent Kits (v3.1 Chemistry Dual Index) (CG000315 Rev C)” (47). “Control agg 3p” dataset is sourced from a healthy donor and consists of 1 full chip (8 channels) run manually and 1 full chip (8 channels) run on the Chromium Connect. Libraries were prepared following either the Chromium Single Cell 3ʹ Reagent Kits v3.1 (CG000204 RevD) or Chromium Next GEM Automated Single Cell 3ʹ Reagent Kits v3.1 (CG000286 RevA) on the Chromium Connect, and were aggregated into this dataset (48).

### Single-cell raw data processing and sample demultiplexing

Using 10x Genomics Cell Ranger v7.1.0 (43), reads were aligned to the human reference genome (10x precompiled reference version GRCh38-2020-A), and gene–cell expression matrices were generated. For hash tag-based sample demultiplexing we devised our own tool called YAHD (Yet Another Hashtag Demultiplexer) as a fast Python tool with minimal dependencies and the possibility for interactive optimization. In particular, we wanted to demultiplex data from partially clogged 10x runs, which was not possible with Cell Ranger. As far as we are aware, such a tool did not exist before.

YAHD is based on a heuristical algorithm which assumes that there are (at least) two peaks in the distribution of counts per hash tag across droplets, corresponding to signal (cell containing droplets) and background (droplets without cells). YAHD also assumes that the hash tags have different affinity and therefore their count distributions must be shifted and scaled before comparing their counts between hash tags.

YAHD works in an iterative way to assign final demultiplexing results to all droplets. It starts with an initial guess on droplets which pass basic QC (at least 100 genes expressed, which are expressed in at least 10 cells, and at least 10 hash tag counts) defined as following: The hash tag counts are scaled by their median per hash tag. Then per droplet an initial “hash tag singlet” demultiplexing annotation is assigned if there is one hash tag which has a higher normalized count than the sum of the normalized counts of the other hash tags. All droplets which do not reach this threshold remain unassigned. Then iteratively using the previous assignments a rescaling for the raw hash tag counts is determined. For every hash tag separately, a characteristic count for hash tag-positive and - negative droplets is estimated (positive: median of the counts over the previously hash tag-positive assigned droplets for that hash tag; negative: the average of those medians for the other hash tags). Then the counts for this hash tag are first scaled by the hash tag-positive count, log-transformed, and then scaled by the log-ratio of hash tag-positive to negative count. After this step, these YAHD-scaled hash tag counts are better comparable between hash tags, with the positive peak centered at 0 and the (highest) negative peak at -1. For all droplets which were assigned hash tag singlets previously, the location of the minimum between the positive and negative peak is found in the common distribution of YAHD-scaled hash tag counts, which is used to define a threshold to call a droplet hash tag positive for the current iteration. The threshold to call a droplet hash tag negative is by default -0.5, i.e. half way between the positive and negative YAHD-scaled count peaks. If a droplet has a YAHD-scaled hash tag count value in between the two thresholds, it is assigned as uncertain for that hash tag. A droplet is assigned “multiplet” if it is hash tag positive for 2 or more hash tags, “empty” if it is hash tag negative for all hash tags, “unassigned” if it is not a multiplet and uncertain for at least one hash tag, and finally assigned singlet for a certain hash tag if it is only positive for that hash tag and negative for all others. This is iterated until consistency, which is defined as the hash tag-positive peak is not larger than 0.1 in YAHD-scaled counts for any of the hash tags. This is usually achieved already after a single iteration.

To compare and evaluate YAHD’s performance we also ran the demultiplexing integrated into Cell Ranger v7.1.0 (36) with the single public parameter min-assignment-confidence at its default value 0.9 as well as vireoSNP v0.5.9 (37) with 2 genotypes (-N 2) based on SNP profiles for each cell generated with cellsnp-lite v1.2.3 (49) (with options -- minMAF 0.1 and -- minCOUNT 20) using their compiled list of 7.4M common variants.

### Single-cell data preprocessing and clustering

Aggregated gene–cell matrices were analyzed with the Scanpy package (50) (https://scanpy.readthedocs.io). Preprocessing included quality-control filtering: cells with less than 1,000 or more than 10,000 total counts, and less than 200 unique transcripts, were removed using pp.filter_cells; genes not detected in at least three cells were excluded with pp.filter_genes. Cells with greater than 25% mitochondrial transcript content were defined as dead in the assessment of cell viability (**Fig. 1b**), while for all other analysis a more stringent threshold of 5% mitochondrial transcript was used as cell filter criterion to ensure that only high quality cells would be analyzed for stress and activation signatures.

Normalization was performed with pp.normalize_total, scaling counts per cell to 10,000. Data were log-transformed with pp.log1p. Highly variable genes were identified using filter_genes_dispersion (parameters: min_mean = 0.0125, max_mean = 3, min_disp = 0.5), resulting in 1,867 highly variable genes.

Principal component analysis (PCA) was performed using tl.pca, and variance explained by each principal component was visualized with pl.pca_variance_ratio. A neighborhood graph was constructed using pp.neighbors (n_neighbors = 10, n_pcs = 40) and embedded with uniform manifold approximation and projection (UMAP) with tl.umap (51). Leiden clustering (tl.leiden) was applied at resolution 1.

Marker gene identification and cluster annotation were performed using CellTypist v1.7.1 (52,53), which employs regularized logistic regression models trained on curated immune reference datasets. Annotations were refined through majority voting across nearest neighbors. Marker genes were identified with tl.rank_genes_groups using the Wilcoxon test, and the top 10 markers per cluster were visualized using sc.pl.rank_genes_groups_matrixplot. Manual curation with known marker genes was also performed.

The computational environment used for annotation included: scanpy==1.9.1, anndata==0.8.0, umap==0.5.3 numpy==1.20.3, scipy==1.9.3, pandas==1.4.3, scikit-learn==1.1.3, statsmodels==0.13.5, python-igraph==0.10.8, louvain==0.8.0, and pynndescent==0.5.12.

### PBMC and pDC enrichment assessment

To assess dead-cell removal by FACS, PBMC viability prior to sorting was measured using the Countess cell counter, and post-processing viability was estimated as the proportion of cells with less than 25% mitochondrial gene content. pDC enrichment was evaluated by comparing pDC ratios across three stages: during FACS as the cell population ratios reported by the FACS machine, input to the 10x channel during loading as calculated based on the sorted cell numbers, and the annotated cell type fractions as calculated in the retrieved scRNA-seq data data.

### Downsampling

To minimize technical variability arising from differences in sequencing depth across samples, per-sample read downsampling was performed based on total UMI counts. For each sample, the mean total counts per cell were first computed, and the sample with the lowest mean value was used as the reference. A downsampling factor was then calculated for each remaining sample as the ratio of the minimum mean total counts to the sample’s mean total counts. Using this factor, raw read counts were proportionally downsampled by sampling from a binomial distribution to achieve comparable UMI counts per cell across all datasets.

### Gene signature analysis

To evaluate cellular stress induced by preprocessing, curated gene sets were obtained for cellular stress response (GO:0033554, 1604 genes post filtering), osmotic stress (GO:0071470, 45 genes post filtering), shear stress (GO:0034405, 15 genes post filtering), and T cell activation (GO:0042110, 495 genes post filtering). Gene lists were filtered for presence across all datasets, and signature scores were computed using tl.score_genes. Mean signature scores for stress responses were compared across Lab-sourced, Clinical, and control datasets with 3′ and 5′ chemistries. T cell activation signatures were computed for “Tcm/Naive helper T cells” and “Tcm/Naive cytotoxic T cells.” Statistical significance was assessed with the Wilcoxon test using the geom_signif function from the R package ggplot2.

### Technical and biological variability assessment

In the Lab-sourced dataset, the experimental setup included two base-line samples per multiplex and together with the channel splitting approach this allowed direct quantification and comparison of biological and technical variability. Biological variability was defined as absolute differences in average stress- and immune-related expression signatures between two participants processed on the same 10x channel (biological replicates). Technical variability was defined as absolute differences for the same participant loaded on two separate 10x channels (technical replicates). The Mann-Whitney U test was used to compare the distributions of expression signature score differences between biological and technical replicates to assess whether observed variability predominantly reflected biological rather than technical sources.

## Data and code availability

Analysis code and data are available on GitHub (https://github.com/KlughammerLab/scRNAseq_mux_FACS). YAHD is available on GitHub (https://github.com/simonwm/yahd). The detailed protocol is available on protocols.io (https://dx.doi.org/10.17504/protocols.io.j8nlk82m5l5r/v1).

## Acknowledgements

We thank all sample donors. We thank Andreas Hauser for maintaining computing infrastructure and Prof. Dr. Max Kaufmann for providing lab access. We gratefully acknowledge LMU Klinikum for providing computing resources on their Clinical Open Research Engine (CORE), the Bioinformatic Core Facility of the Biomedical Center Munich for providing computing resources on their HPC system, the Core Facility Flow Cytometry (CFFlowCyt) at the Biomedical Center Munich for their support with cell sorting device usage, and the LAFUGA technology platform at the Gene Center for high-throughput sequencing.

## Author contributions

S.W.M. and J.K. designed the study with input from M.M. and T.J.S.R. M.M. and T.J.S.R. developed the protocol with input from A.K., M.K., and J.K.. A.B., P.B., M.S. and H.W. recruited participants for the exercise study (Lab-sourced dataset). C.W. and M.L-K. recruited patients for the interferonopathy study (Clinical dataset). M.M., T.J.S.R., A.B., P.B., M.S. and C.W. processed blood samples. M.M. and T.J.S.R. generated the scRNA-seq data with support and input from A.E., A.K., V.P., and from G.D., who also assisted on the initial transference of the protocol into its online published format. S.W.M. developed YAHD and processed the raw data. M.M. performed validation analysis with support from A.E., T.J.S.R., J.K. and S.W.M.. M.M., T.J.S.R.,S.W.M. and J.K. wrote the manuscript with input from all co-authors. All authors read and approved the final manuscript.

---

**Supplemental Figure S1.**
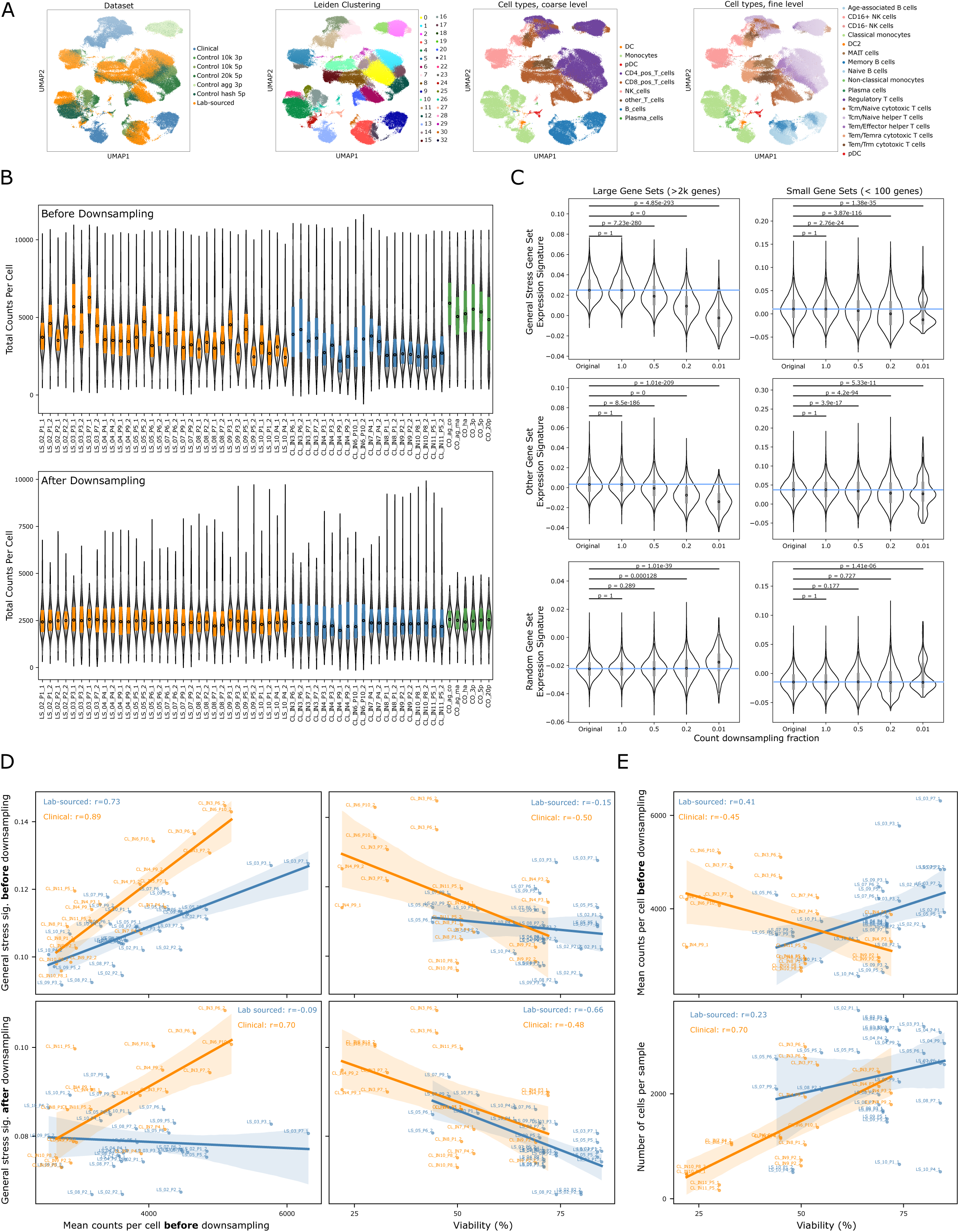
Dataset overview and quality control. (A) UMAP representation of all datasets post quality control and filtering, displaying cell clustering and annotations at both coarse and fine cell type levels. (B) Violin and box plots displaying sequencing depth represented by UMI counts per cell and its normalization through downsampling across samples. Top: Distribution of UMI counts per cell across the Lab-sourced, Clinical, and control PBMC datasets prior to downsampling, illustrating substantial variation due to differences in sequencing depth. Bottom: Distributions after downsampling, demonstrating normalization of UMI counts per cell distributions across samples. (C) Violin and box plots showing the distribution of expression signature scores per cell, for different degrees of UMI count downsampling (1.0, 0.5, 0.2, 0.01 of original total UMI count) of a 10x control dataset (Control 10k 5p). Results for three types of gene sets (GO general stress gene set, another GO gene set of similar size, set of randomly selected genes) in a large (> 2000 genes) and a small (<100 genes) version are displayed. Blue lines indicate the median for the original, non-downsampled data for comparison. (D) Scatter plots showing the relationships between average UMI counts per cell or sample viability (prior to FACS) and average general stress expression signature scores before (top) and after (bottom) downsampling. (E) Scatter plots showing the relationships between sample viability (prior to FACS) and average UMI counts per cell before downsampling (top) or number of cells per sample (bottom).

